# Characterization of the mitochondrial proteome in the ctenophore *Mnemiopsis leidyi* using MitoPredictor

**DOI:** 10.1101/2023.08.23.554511

**Authors:** Viraj Muthye, Dennis V. Lavrov

**Author notes:** **Corresponding author:** Dennis V. Lavrov **Email:**.

## Abstract

Mitochondrial proteomes have been experimentally characterized for only a handful of animal species. However, the increasing availability of genomic and transcriptomic data allows one to infer mitochondrial proteins using computational tools. MitoPredictor is a novel random-forest classifier, which utilizes orthology-search, mitochondrial targeting signal (MTS) identification, and protein domain content to infer mitochondrial proteins in animals. MitoPredictor’s output also includes an easy-to-use R Shiny applet for the visualization and analysis of the results. In this article, we provide a guide for predicting and analyzing the mitochondrial proteome of the ctenophore *Mnemiopsis leidyi* using MitoPredictor.

## 1. Introduction

Best known for their role in energy metabolism ^1^, mitochondria are involved in a multitude of cellular processes, including Fe/S cluster biosynthesis ^2^, amino-acid metabolism ^3^, lipid metabolism ^4^, apoptosis ^5,6^, and cellular signaling ^7^. The diverse mitochondrial functions require more than a thousand proteins, the vast majority of which are encoded in the nuclear genome and transported into the mitochondria via several import pathways ^8,9^. Collectively, these nuclear-encoded mitochondrial proteins constitute the bulk of the “mitochondrial proteome,” an inclusive collection of proteins with at least some mitochondrial functions. A few additional proteins are encoded in the mitochondrial genome – mtDNA, which is replicated, repaired, and expressed mostly independently from the nuclear genome ^8^. Information on the mitochondrial proteome should be useful for understanding the functional repertoire of mitochondria. However, our ability to distinguish mitochondrial proteins remains limited.

Mitochondrial proteomes have been experimentally characterized for a few animals ^10–13^, plants ^14–17^, fungi ^18^, and protists ^19^. In animals, experimental characterization of the mitochondrial proteome is limited to model animal species, including *Homo sapiens* ^10,11^, *Mus musculus* ^10,11^, *Caenorhabditis elegans* ^12^, and *Drosophila melanogaster* ^13^. Most animal phyla – including the four non-bilaterian phyla (Ctenophora, Porifera, Placozoa, and Cnidaria) – lack experimental mitochondrial proteomic data. This is unfortunate, because non-bilaterian taxa represent most of the oldest lineages in the animal phylogenetic tree and thus are essential for understanding the full extent of animal diversity. They also display some of the most unusual mitochondrial genome organization among animals ^20^.

Computational techniques provide a complementary approach to experimental characterization of mitochondrial proteomes. They attempt to infer mitochondrial proteins from genomic and transcriptomic data, which are becoming increasingly available even for non-bilaterian taxa. Several subcellular localization predictors have been developed and can be divided into four broad categories:

- **Homology-based predictors** (*e.g*. OrthoFinder ^21^, InParanoid ^22^, Proteinortho ^23^): One way to predict mitochondrial proteins is by finding orthologs in experimentally characterized mitochondrial proteomes. The advantage of this approach is that it can detect mitochondrial proteins, which lack identifiable mitochondrial targeting signals (MTS), as is the case for most mitochondrial inner-membrane proteins and all outer-membrane proteins. Its disadvantage is that it would fail to identify novel proteins without known orthologs and might be misled by instances of protein subcellular re-localization.
- **N-terminus Mitochondrial Targeting Signal (MTS) predictors** (*e.g*. TargetP ^24^, MitoFates ^25^): Another common method for inferring mitochondrial proteins is by searching for the mitochondria targeting signal (MTS) at the protein’s N-terminus. MTS is a short 10-90 amino-acid sequence, enriched in positively charged amino-acids that directs proteins towards the mitochondria. Once inside the mitochondrial matrix, the MTS is cleaved, and the mature protein is formed ^25^. Because MTS predictors do not rely on homology, they can identify novel mitochondrial proteins. However, these predictors tend to suffer from a high false-positive rate and would not identify mitochondrial proteins possessing a non-canonical MTS.
- **Full protein sequence feature-based predictors** (*e.g*. CELLO2.5 ^26^, ngLOC ^27^, SubMitoPred ^28^): Some predictors utilize features extracted from the whole protein sequences, like amino-acid composition, protein domain composition, hydrophobicity, *etc*. to infer mitochondrial localization. CELLO uses n-peptide composition, partitioned amino-acid composition, g-gap dipeptide composition, and local amino acid composition; ngLOC uses the density-distribution of peptide sequences of a fixed length (*n-gram*s); while SubMitoPred – protein domain composition and several additional features extracted from the input protein sequences.
- **Ensemble predictors** (*e.g*. MitoPredictor ^29^, SubCons ^30^): Ensemble methods use machine-learning algorithms to integrate multiple sources of information for predicting mitochondrial proteomes. MitoPredictor is a random forest-based predictor, which utilizes 1)orthology, 2) MTS prediction and 3) protein domain information. SubCons uses a random forest classifier to integrate results from four independent programs: CELLO2.5, SherLoc2 ^31^, LocTree2 ^32^, and MultiLoc2 ^33^.

While each of the four categories of predictors have their advantages and disadvantages, ensemble methods tend to outperform other methods because they integrate information from multiple sources. In this article, we provide a guide for predicting and analyzing the mitochondrial proteome in the ctenophore *Mnemiopsis leidyi* using our recently developed ensemble predictor MitoPredictor. MitoPredictor provides several advantages over other ensemble predictors and has been shown to outperform SubCons, a widely-used ensemble protein predictor of animal protein subcellular localization. In addition to running the analysis, MitoPredictor provides an easy-to-use R Shiny applet for visualization and exploration of the results. The applet also includes information on existing experimentally-characterized mitochondrial proteomes from human, mouse, *C. elegans*, and *D. melanogaster*.

Ctenophores are the prime candidate for characterization of their mitochondrial proteome. They constitute an ancient lineage in animal evolution and, possibly, the sister group to the rest of the animals ^34^. They also have some of the most unusual mitochondrial genomes among animals^35^. While nuclear and mitochondrial genomes from multiple ctenophores have been sequenced (Tables 1 and 2), no mitochondrial proteomes have been experimentally characterized from this group.

**Table 1.** Publicly-available nuclear genomes from Ctenophora.

| Species | GenBank ID |
| --- | --- |
| *Pleurobrachia bachei* | GCA_000695325.1<sup>38</sup> |
| *Bolinopsis microptera* | GCA_026151205.1<sup>34,39</sup> |
| *Hormiphora californensis* | GCA_020137815.1<sup>40</sup> |
| *Beroe forskalii* | GCA_011033025.1<sup>41</sup> |
| *Beroe ovata* | GCA_946803715.1 |
| *Mnemiopsis leidyi* | GCA_000226015.1<sup>36</sup> |

**Table 2.** Publicly-available mitochondrial genomes from Ctenophora.

| Species | GenBank ID |
| --- | --- |
| *Beroe forskalii* isolate Bf201311 | MG655624.1<sup>41</sup> |
| *Beroe forskalii* isolate Bf201606 | MG655623.1<sup>41</sup> |
| *Beroe forskalii* isolate Bf201706 | NC_038065.1<sup>41</sup> |
| *Hormiphora californensis* isolate Hc2 | MN544301.1<sup>40</sup> |
| *Hormiphora californensis* isolate Hc1 | NC_045864.1<sup>40</sup> |
| *Hormiphora californensis* isolate 20161213-T1 | CM035363.1<sup>40</sup> |
| *Pleurobrachia bachei* | JN392469.1<sup>42</sup> |
| *Beroe cucumis* | NC_045305.1<sup>43</sup> |
| *Bolinopsis microptera* isolate Bmic1 | CM047416.1<sup>39</sup> |
| *Mnemiopsis leidyi* | NC_016117.1<sup>35</sup> |

Here, we used MitoPredictor to predict and analyze the mitochondrial proteome of *M. leidyi*. This ctenophore has one of the smallest mitochondrial genomes in animals (∼10 kb) that lacks all tRNA genes as well as *atp6* and *atp8* ^35^. Additionally, mtDNA-encoded ribosomal RNA molecules exhibit extremely reduced primary and secondary structures. *M. leidyi* mtDNA also exhibits a very high rate of mitochondrial sequence evolution. It is likely that some of these changes would necessitate changes in its mitochondrial proteome. For example, we have shown that mt-ribosomal proteins encoded in the nucleus have undergone expansions, possibly to compensate for the reduction in the rRNA structures ^35^. While *M. leidyi* lacks an experimentally-characterized mitochondrial proteome, there is a high-quality nuclear genome, which is deposited at the *Mnemiopsis* Genome Project Portal (https://research.nhgri.nih.gov/mnemiopsis/) ^36,37^.

## 2. Materials and methods

### 2.1 Software required

1. MitoPredictor: https://github.com/virajmuthye/mitopredictor.
2. Proteinortho v5.16b: https://www.bioinf.uni-leipzig.de/Software/proteinortho/.
3. TargetP v1.1: http://www.cbs.dtu.dk/services/TargetP-1.1/index.php
4. MitoFates: http://mitf.cbrc.jp/MitoFates/cgi-bin/top.cgi
5. CD-HIT ^44^: http://weizhongli-lab.org/cd-hit/
6. R (>= 3.0.2): https://www.r-project.org/
7. R studio: https://rstudio.com/
8. Perl: https://www.perl.org/get.html

### 2.2 Databases required

1. Pfam-A.hmm: ftp://ftp.ebi.ac.uk/pub/databases/Pfam/releases/ (Select latest release)
2. *Mnemiopsis leidyi* protein models from the *Mnemiopsis* Genome Portal: https://research.nhgri.nih.gov/mnemiopsis/download/download.cgi?dl=proteome. There are two categories of protein models in the portal: *Filtered* and *Unfiltered*. Filtered proteins models were used for the analysis.

### 2.3 Identifying mitochondrial proteins using MitoPredictor

MitoPredictor consists of seven BASH scripts and an R script that process input data and invoke the computational tools listed above (Section 2.1) to predict mitochondrial proteins. Each step of MitoPredictor requires only a single command: *e.g*., *step1_prep.bash*, which would run this script using default settings. The information on changing the default settings are specified within each individual script.

The input file for MitoPredictor is a set of amino-acid sequences from the query species in a FASTA format. The FASTA format consists of at least two lines for each entry: 1) the first line beginning with the “>” symbol and contains the name of and some additional (optional) information about the entry and 2) the amino-acid sequence. An example of a protein entry in the FASTA format is given below:

~~~
>sp|Q8ITI5|FOXG1_MNELE Forkhead box protein G1 OS=Mnemiopsis leidyi OX=27923 GN=FOXG1 PE=2 SV=1
MVVTTATKPHPFSIENILKSASPKPQKPLFSYNALIAMAISQSPLKKLTLSEIYDFIIET
FPYYRDNKKGWQNSIRHNLSLNKCFVKVPRHYNDPGKGNYWMLNPNSDEVFIGGKLRRRP
GQNGGSLESYMHLKTRTSPYQRGDTVCKRDRVVYLSNSGAGNCQFYQPVPCSSPTAMLSR
SSLIVQTSPTTLTIPHHPPQNYIQTLSPNVSPVRVGNQTSPRQSALPSSSLPLLPSPPSL
PSKLSSPPLPSLSTNLPSPPLDTGDILLSPFLRQTVDSTSQNFLEHMIRIREQVQRSGHL
ALAQSRAVVYQPIPRKSL
~~~

The software and databases listed above need to be downloaded and installed prior to running MitoPredictor. The input proteome, *i.e*., protein models from the *Mnemiopsis* Genome Project Portal, should be placed in the MitoPredictor directory. MitoPredictor has been developed and tested on a Linux cluster but should be transferable to other systems.

#### 1. Step I: Pre-processing *Mnemiopsis* proteins

This step is run as *step1_prep.bash*. (See **Note 1** and **Note 2** prior to running Step I)

- MitoPredictor will identify the input file which should have the “.fasta” file extension.
- Next, all potential fragments, *i.e*., proteins below 100 amino acids in length, are removed.
- CD-HIT clusters the remaining proteins at 98% similarity. This cut-off can be changed in the script “prep.bash” inside the prep folder. These proteins are used as input for step II (orthology) and step IV (protein domain analysis).
- All proteins without methionine at position 1 are removed. The remaining proteins are used as an input for step III (MTS prediction). Removal of potential fragments that are missing a complete N-terminus is aimed at reducing the false-positive rate in step II.

#### 2. Step II: Orthology analysis

This step is run as *step2_ortho.bash*.

- The script invokes Proteinortho, which identifies groups of orthologous proteins (OGs) in the reference species (human, mouse, *C. elegans, D. melanogaster*, and *S. cerevisiae)* and the query species (*M. leidyi*). Default settings are used for Proteinortho: e-value: 1e^-05^, identity: 25%, minimum coverage of best BLAST hit alignments: 50%, algebraic connectivity: 0.1, and purity: 0.1 (these values can be changed in the “*run_proteinortho.bash*” script in the “orthology” folder). A four-letter abbreviation is used to denote each of the five reference species (Table 3).
- One feature is extracted in this step: OrthoScore (OS). For each *M. leidyi* protein, the OS is 1 if an ortholog is found in at least one reference mitochondrial proteome, and 0 if not.

**Table 3.** The four-letter abbreviations that were used to denote each reference species in MitoPredictor.

| Species | Abbreviation |
| --- | --- |
| *Homo sapiens* | hsap |
| *Mus musculus* | mmus |
| *Caenorhabditis elegans* | cele |
| *Drosophila melanogaster* | dmel |
| *Saccharomyces cerevisiae* | scer |

#### Step III: MTS analysis

This step is run as *step3_mts.bash*.

- TargetP and MitoFates predict MTS in *M. leidyi* proteins. Default settings are used for TargetP and MitoFates but can be changed in the “*runTargetP.bash*” and “*runMitoFates.bash*” scripts in the “mts” folder.
- For each *M. leidyi* protein, five features are extracted from the results generated by the two programs:
  i. From TargetP: mTP (probability that the protein possesses an MTS), sTP (probability that the protein possesses a signal peptide), other (probability that the protein localized to any other location)
  ii. From MitoFates: mfprob (probability that the protein possesses an MTS) and net-charge.

#### 3. Step IV: Protein domain analysis

his step is run as *step4_domain.bash*.

- Protein domains are identified in *M. leidyi* proteins using the Perl script *pfam_scan.pl* (ftp://ftp.ebi.ac.uk/pub/databases/Pfam/Tools/) and the Pfam-A.hmm libraries.
- Based on the protein domain content of each *M. leidyi* protein, a domain score (DS) is assigned to that protein. The DS ranges from 0-1, where a score of 0 indicates that the protein domain exists only in non-mitochondrial proteins in all reference animal species, while a score of 1 indicates that the protein domain has been found only in mitochondrial proteins in the majority of the reference species.

#### 4. Step V: Prediction of mitochondrial proteins using a random forest classifier This step is run as *step5_identify_and_concat.bash*

- In this step, a random forest classifier is used with the features extracted in steps II, III and IV to predict mitochondrial proteins. The classifier is implemented in R (‘*rf.R*’), and uses the R package “randomForest” (https://www.stat.berkeley.edu/~breiman/RandomForests/).

#### 5. Step VI: Make basic statistics of the predicted mitochondrial proteome This step is run as *step6_make_stats.bash*

- MitoPredictor generates basic statistics of the predicted *M. leidyi* mitochondrial proteome, and data for the R Shiny applet (Figure 2).
- Step VI generates two primary outputs:
  i. The “FINAL.MATRIX” file: This file serves as an input for the R Shiny applet “mito_app.R”. The R shiny applet can be accessed in R Studio via the “Run App” button, or by giving the R command runApp(“[directory where the app files are stored]”). The applet can be used to visualize, analyze, and download the mitochondrial proteins from *M. leidyi* and the reference species.
  ii. The “stats” folder: This folder contains information on the predicted *M. leidyi* mitochondrial proteome. More information is provided in Section 2.5.

**Figure 1.**
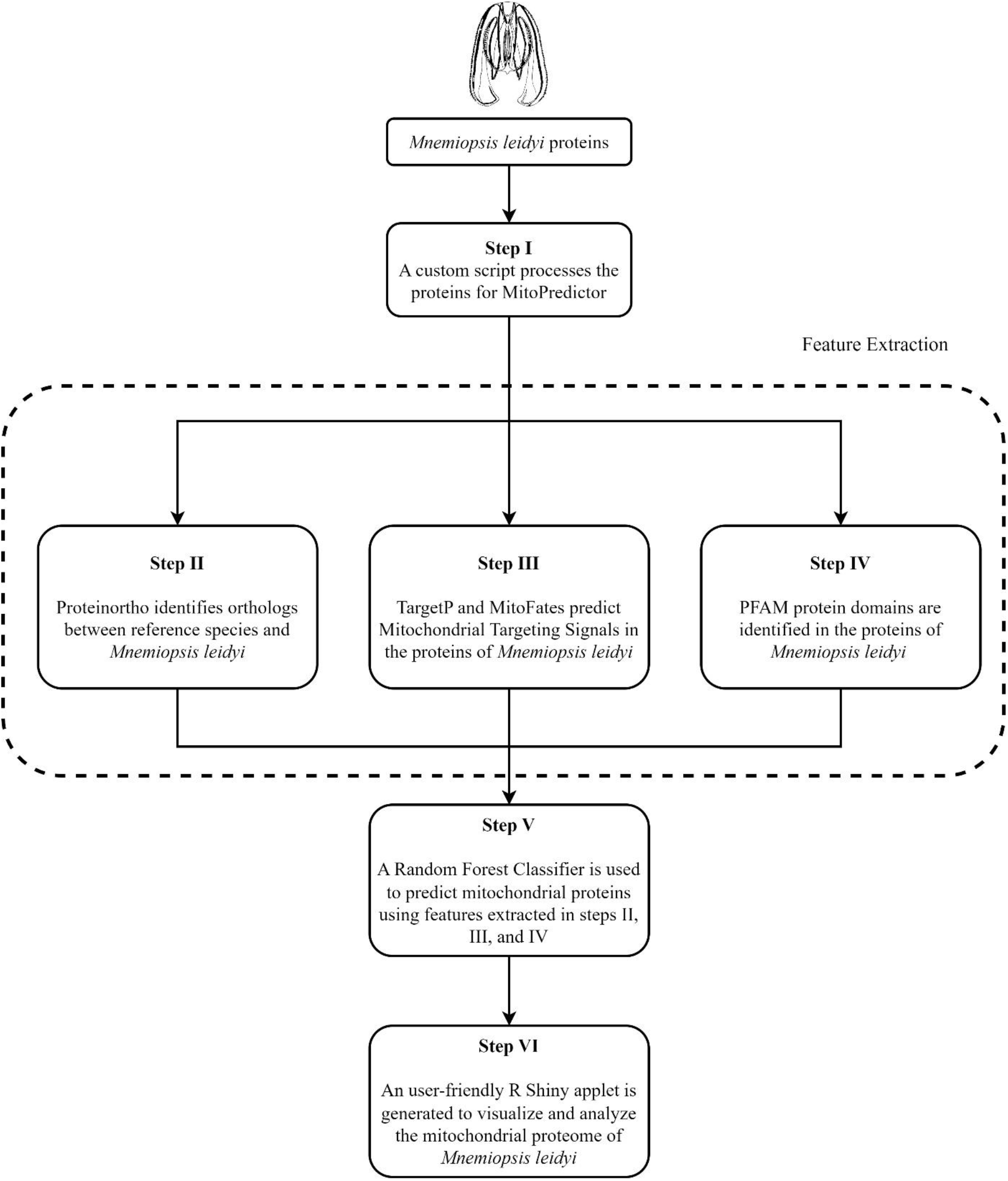
Flowchart outlining the basic steps used by MitoPredictor for predicting mitochondrial proteins in the ctenophore *Mnemiopsis leidyi*. The silhouette of *M. leidyi* was taken from PhyloPic V2 (www.phylopic.org).

**Figure 2.**
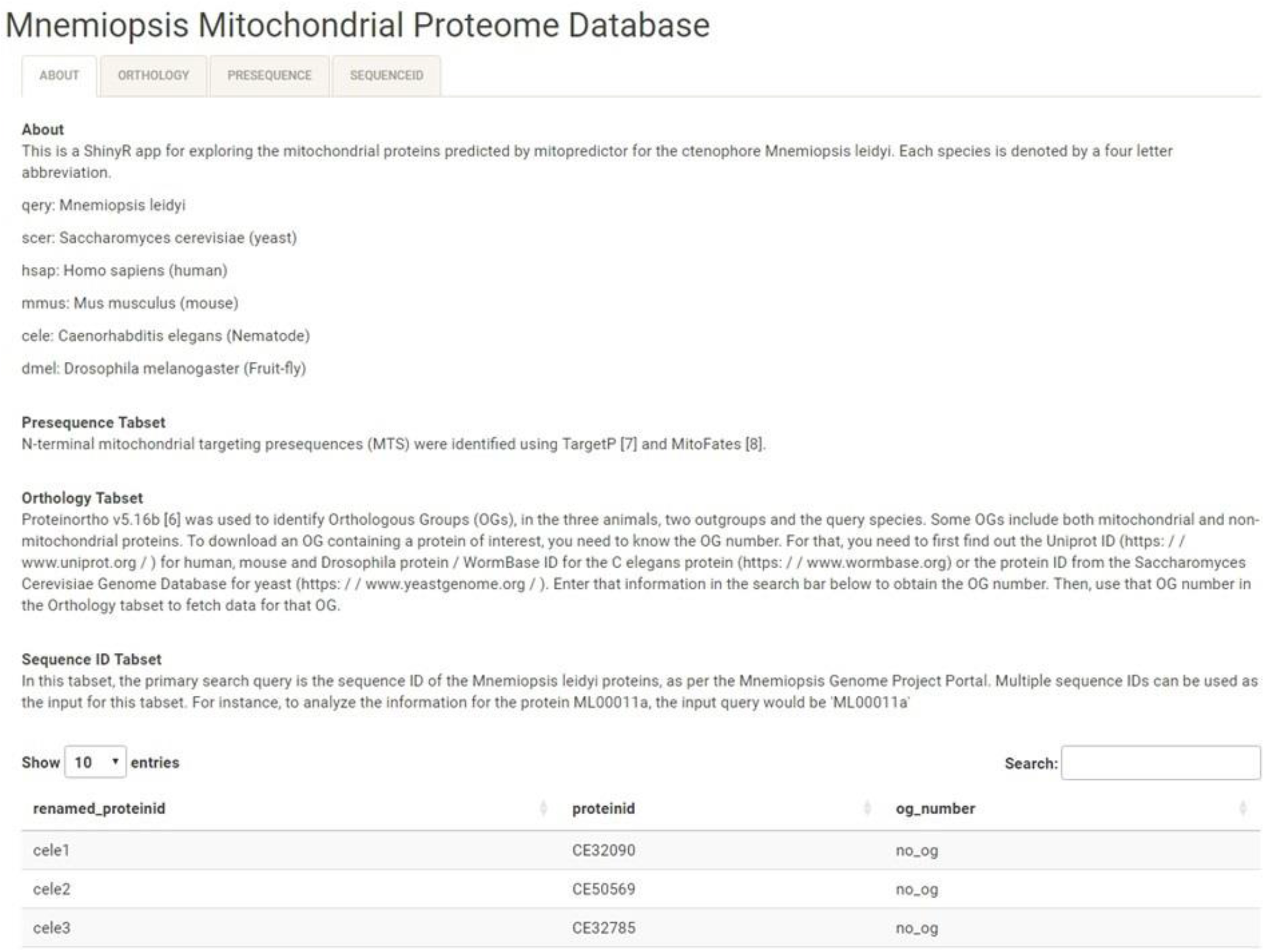
The About tabset of the R Shiny applet generated by MitoPredictor. The applet consists of four tabsets: About, Orthology, Presequence, and Sequence ID.

#### 6. Step VII: Clean intermediate files and prepare MitoPredictor for the next analysis This step is run as *step7_clean.bash*

This step removes intermediate and result files generated during the MitoPredictor run in preparation for the next analysis. It is important to save the results of the run (the “FINAL.MATRIX” file, “mito_app.R” file and the “stats” folder) before running this step.

### 2.4 R Shiny applet

The R Shiny applet allows for the visualization and analysis of the predicted mitochondrial proteome. It is designed to facilitate comparative analysis of the predicted mitochondrial proteome (in the present case, the *M. leidyi* mitochondrial proteome) with the reference mitochondrial proteomes. The “FINAL.MATRIX” file generated in MitoPredictor Step 6 above serves as the input for the applet. The R Shiny applet is organized into four tabsets: 1] About, 2] Orthology, 3] Presequence, and 4] SequenceID (Figure 2). The “About” tabset provides general information and description of other tabsets. Here, we describe the remaining three tabsets:

- **Orthology**: The orthology tabset allows users to analyze results of orthology analysis from Proteinortho. Users can download information on a particular orthology-group (OG) of interest. The primary search query in this tabset is the OG number assigned during analysis. The OG number can be fetched by using either the Uniprot ID for human, mouse, and *D. melanogaster* or the *C. elegans* protein ID from WormBase.
- **Presequence**: This tabset includes MTS-prediction results from TargetP and MitoFates for all the proteins from reference and query species.
- **SequenceID**: This tabset is meant for visualization and analysis of query proteins. In this case, users can extract information for a list of *M. leidyi* proteins. The input for this tabset is the protein ID assigned to the protein on the *Mnemiopsis* Genome Project Portal (*e.g*., “ML000111a”).

### 2.5 Using the information in the “stats” folder to analyze mitochondrial proteomes

The “stats” folder includes information on the predicted mitochondrial proteome. Here, we describe how files in the “stats” folder can be used for the functional annotation of mitochondrial proteins and for the further analysis of the predicted mitochondrial proteome.

- **Stats.txt**: The file “Stats.txt” provides a brief overview of the predicted mitochondrial proteome (Table 4).

**Table 4.**
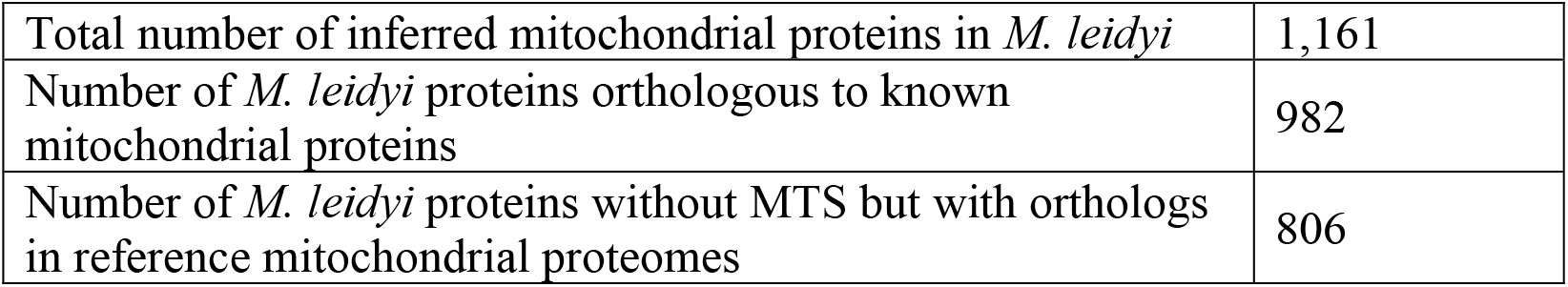
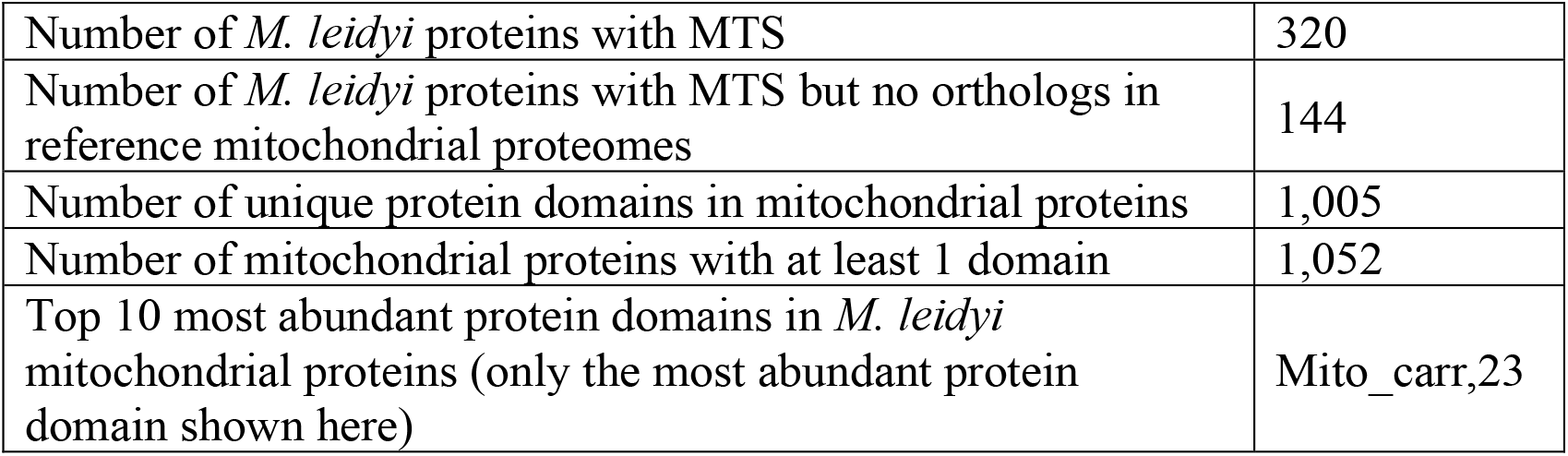
Statistics for the predicted mitochondrial proteomes thar are listed in the file “Stats.txt”.

Based on this file, we can see that MitoPredictor predicted 1,161 mitochondrial proteins in *M. leidyi*. Interestingly, the number of *M. leidyi* proteins possessing a recognizable MTS was low (320), with most of the mitochondrial proteins predicted primarily by the results of the orthology search (982). Among *M. leidyi* proteins with detected MTS, nearly half (144/320) had no ortholog in any of the reference species. 1,005 protein domains were identified in the predicted mitochondrial proteome of *M. leidyi*. The mitochondrial carrier domain (“Mito_carr”) was the most abundant protein domain, present in 23 mitochondrial proteins.

- **species_specific.matrix**: The file “species_specific.matrix” provides information on *M. leidyi* proteins predicted to possess an MTS with no orthologs in reference mt-proteomes. These represent potential *M. leidyi-*specific or ctenophore-specific mitochondrial proteins (but see **Note 5**). The protein domain composition of these proteins, provided in the file, can be used to predict their potential function. The FASTA sequences of these proteins can be found in the file “query_species_specific.fasta”. This file can be uploaded in tools like PANNZER ^45^ (see **Note 6)** or dcGO ^46^ for functional annotation.

The *Mnemiopsis* Genome Project Portal also has several tools for functional analysis of *M. leidyi* proteins. For example, to identify the function of the species-specific *M. leidyi* protein “ML189344a”, a user can enter this name in the “View Gene Page” at the *Mnemiopsis* Genome Project Portal (https://research.nhgri.nih.gov/mnemiopsis/genes/genewiki.cgi). The “View Gene Page” includes information such as functional annotation by Argot2 and Blast2GO and BLAST searches against various databases.

- **domain_count.matrix**: Protein domain composition is known to be useful in functional annotation of proteins and predicting subcellular localization. This file contains a list of protein domains predicted to be present in at least one *M. leidyi* protein and the number of mitochondrial proteins containing each protein domain. The file “Stats.txt” contains a list of the top five most abundant protein domains in the predicted mitochondrial proteome. Here, we see that “Mito_carr” was the most abundant protein domain, found in 23 predicted mitochondrial proteins.
- **<Species>_query_mitoorthologs.txt:** For each reference species, the “stats” folder contains a list of mitochondrial proteins with and without orthologs in *M. leidyi*. For instance, the file “hsap_query_mitoorthologs.txt” contains a list of human mitochondrial proteins which possess orthologs in *M. leidyi*. This file can be used as input for several functional annotation tools, like ConsensusPathDB ^47^, PantherDB ^48,49^, and StringDB ^50^. Below, we provide a list of resources which can be used to analyze these lists provided by MitoPredictor as output (Table 5).

**Table 5.** List of resources for functional annotation of mitochondrial proteins. The gene-lists generated by the MitoPredictor output can be used directly as input for these resources.

| Species | Resources |
| --- | --- |
| Human | PantherDB, ConsensusPathDB, StringDB, UniProt |
| Mouse | PantherDB, ConsensusPathDB, StringDB, UniProt |
| *C. elegans* | PantherDB, StringDB, WormBase <sup>51</sup> |
| *D. melanogaster* | PantherDB, StringDB, FlyMine <sup>52</sup> |

## 3. Notes

1. The input file for Step I requires a “.fasta” file extension. Step I (processing) is *not* a mandatory step. The file “prep.bash” in the “prep” folder provides information on how to skip step I. However, the presence of a large number of C-terminus fragments will result in errors in MTS prediction by TargetP and MitoFates, which require complete N-terminus for accurate prediction of MTS. Therefore, we highly recommend not skipping step 1.
2. The CD-HIT cut-off used in this analysis is 98%. This cut-off can be changed in the file “prep.bash”. While changing the cut-off used in CD-HIT in this step, it is important to make the appropriate corresponding change to the choice of word size (Table 6).
3. While Proteinortho was selected as the tool to infer orthology, there have been several orthology predictors that have been developed since then, like OrthoFinder. However, MitoPredictor has not been tested using OrthoFinder or any other software.
4. Steps II, III, and IV can be run in parallel to reduce the runtime of MitoPredictor. Step V, the Random Forest classifier requires the features extracted from steps II-IV and should be run only after all the previous steps are finished.
5. In the output of MitoPredictor, a “species-specific” mitochondrial protein is a protein that lacks orthologs in the five reference proteomes (human, mouse, *C. elegans, D. melanogaster*, and yeast). However, some of these “species-specific” mitochondrial proteins can be orthologous to mitochondrial or non-mitochondrial proteins in reference species, which are missed by the orthology search applied by Proteinortho. Some “species-specific” mitochondrial proteins could also be orthologs of mitochondrial or non-mitochondrial proteins from species not included in MitoPredictor. Finally, it is also possible that some of these “species-specific” proteins are false-positives of MTS prediction. We recommend running Protein BLAST (BLASTP) on “query_species_specific.fasta” after running MitoPredictor. BLASTP can be downloaded and run locally or accessed at (https://blast.ncbi.nlm.nih.gov/Blast.cgi). In *M. leidyi*, 23/144 species-specific proteins produced no hits in the NR database (non-redundant protein sequences), while 121 showed similarity to some known proteins in other species. It is important to note that a BLASTP hit does not necessarily imply the presence of a homolog.
6. Pannzer can be accessed via the Pannzer2 web server (http://ekhidna2.biocenter.helsinki.fi/sanspanz/). The webserver accepts a file of protein sequences in FASTA format as an input. GO predictions from four predictors (ARGOT, RM3, JAC, and HYGE) are provided by Pannzer2. ARGOT has been shown to be the top performing predictor.
7. In case users have a transcriptome for the species of interest, TransDecoder can be used to predict protein models prior to running MitoPredictor (https://github.com/TransDecoder/TransDecoder/wiki). TransDecoder, along with its instructions for usage, can be downloaded from https://github.com/TransDecoder/TransDecoder/wiki. TransDecoder generates several output files. The output file with the “.transdecoder.pep” file extension should be used as the input file for MitoPredictor. We also recommend using CD-HIT to cluster the predicted proteins at 100% similarity.

**Table 6.** The word-size for threshold used, as recommended by the CD-HIT manual.

| Word-size (-n) | Threshold (cut-off) |
| --- | --- |
| 5 | 0.7 - 1 |
| 4 | 0.6 – 0.7 |
| 3 | 0.5 – 0.6 |
| 2 | 0.4 – 0.5 |

## 4. Analysis of the predicted *M. leidyi* mitochondrial proteome

### 1. Conservation and loss of common animal mitochondrial proteins

Any investigation into a novel mitochondrial proteome should infer potential losses and gains of function. To study the loss of mitochondrial function, one can consider the orthologous groups (OGs) that contain proteins from the reference species, but not the species of interest (*M. leidyi*). In our study, we identified 255 OGs containing at least one mitochondrial protein from at least three out of the four reference animal species and yeast. 179/255 of them included at least one predicted mitochondrial protein from *M. leidyi*. However, 76 did not include any *M. leidyi* ortholog, suggesting a potential loss in this species (Figure 3A). Next, we analyzed the function of the human mitochondrial proteins contained in these 76 OGs using PantherDB. The top five enriched GO Biological Process terms were “tRNA aminoacylation for mitochondrial protein translation (GO:0070127)”, “mitochondrial translational termination (GO:0070126)”, “asparaginyl-tRNA aminoacylation (GO:0006421)”, and “[4Fe-4S] cluster assembly (GO:0044572)” (Figure 3C). This observation is consistent with our previous study that found that the loss mitochondrial tRNA genes in *M. leidyi* co-occurred with the loss of most mitochondrial aminoacyl tRNA synthetases ^53^.

**Figure 3.**
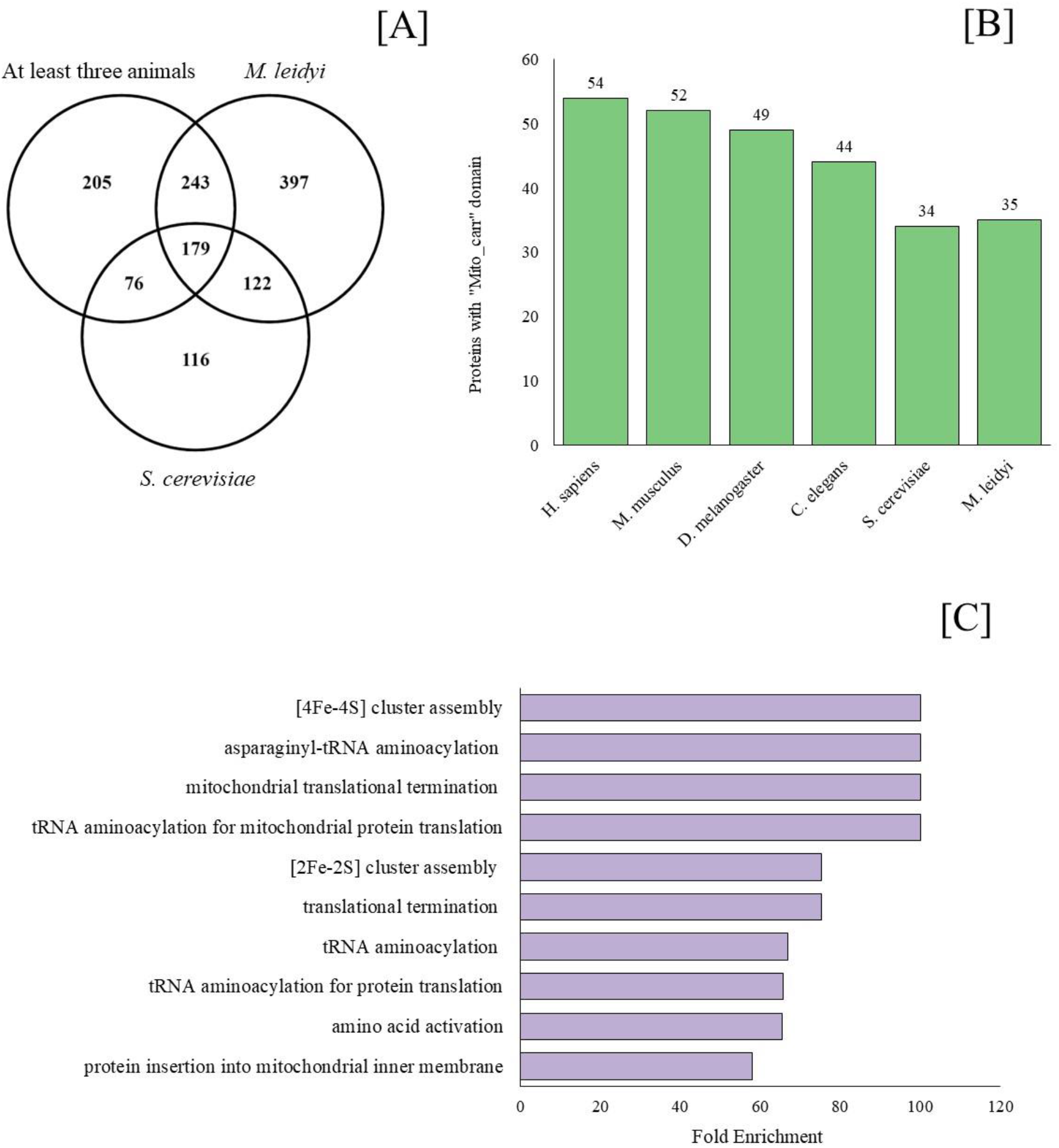
**A]**, Venn diagram of OGs (groups of orthologous proteins predicted by Proteinortho) containing mitochondrial proteins from at least three out of the four reference animal species, *M. leidyi*, and yeast **B]**, The number of proteins with “Mito_carr” protein domains, **C]** Functional annotation of proteins from 76 OGs that contain mitochondrial proteins from the majority of reference animals and yeast but not from *M leidyi*. Functional annotation was performed using the human proteins from these 76 OGs in PantherDB.

### 2. Analysis of mitochondrial carrier proteins in *M. leidyi*

Mitochondria can be subdivided into four main compartments: mitochondrial outer membrane, mitochondrial inner membrane, mitochondrial intermembrane space, and mitochondrial matrix. The mitochondrial outer membrane is permeable to certain solutes (molecular mass less than 5 kDa). However, the mitochondrial inner membrane is impermeable to most molecules. Mitochondrial carrier proteins are hydrophobic inner membrane proteins involved in the transport of specific substrates, like ADP, ATP, NAD+, succinate, fumarase, malate, and phosphate, across the mitochondrial inner membrane that are present in all major eukaryotic groups ^54,55^.

In the four reference animal species, the number of mitochondrial carrier proteins ranges from 44 in *C. elegans* to 54 in human. We found that the number of mitochondrial carrier proteins in *M. leidyi* (35) was lower than those in the bilaterian animals, but similar to that in yeast (Figure 3B). Mitochondrial carrier proteins not identified in *M. leidyi* included the tricarboxylate carrier protein, mitochondrial uncoupling protein 4, mitochondrial brown fat uncoupling protein 1, and mitochondrial ornithine transporter 2. Further analysis would be required to identify whether and how these substrates are imported into the mitochondria in *M. leidyi*.

### 3. Species-specific mitochondrial proteins in *M. leidyi*

Out of 320 *M. leidyi* proteins predicted to possess an MTS, 144 lacked an ortholog in any reference mitochondrial proteomes. 63 of these 144 proteins possessed at least one predicted protein domain. These “species-specific” proteins can represent instances of neolocalization of cytosolic proteins, known mitochondrial proteins with high rates of sequence evolution, or novel mitochondrial proteins. Mitochondrial neolocalized proteins, *i.e*., predicted mitochondrial proteins in *M. leidyi* with only non-mitochondrial orthologs in reference species are the easiest to detect and analyze. We identified several such proteins in *M. leidyi*, including five cytosolic ribosomal proteins, DNA methyltransferase 1-associated protein 1 (DMAP1), and NAD-dependent protein deacetylase sirtuin-2 (SIR2). All these proteins in *M. leidyi* possessed an MTS sequence predicted at high probability.

Additionally, we detected a *M. leidyi* protein containing an “MFS_1” domain, belonging to the Major Facilitator Superfamily (MFS), with no orthologs in any reference proteomes. The Major Facilitator Superfamily is the largest group of secondary membrane transporters that is responsible for the transport of a wide range of substrates across various membranes ^56,57^. BLASTP identified potential homologs of the *M. leidyi* MFS protein in several prokaryotes, but only three eukaryote species. Interestingly, the majority of the eukaryotic sequences possessed an MTS as predicted by MitoFates (4/6 sequences). Experimental characterization of these predicted species-specific mitochondrial proteins needs to be performed to confirm their mitochondrial localization and function.

## 5. Conclusion

In this article, we present a guide for the prediction and analysis of nuclear-encoded mitochondrial proteins (mitochondrial proteome) in the ctenophore *M. leidyi* with MitoPredictor. MitoPredictor, a random forest-based classifier, uses orthology search, MTS prediction, and protein domain information for inferring mitochondrial proteins in animals. In addition, MitoPredictor provides an easy-to-use R Shiny applet for the visualization and analysis of the results. The output of MitoPredictor is also well-suited for further functional analysis using several existing tools, like PantherDB and ConsensusPathDB. MitoPredictor identified 1,162 mitochondrial proteins in *M. leidyi*. 156 of these predicted mitochondrial proteins did not possess an ortholog in the five reference species included in MitoPredictor. Several well-conserved mitochondrial proteins were not identified in *M. leidyi*, including multiple mitochondrial aminoacyl tRNA synthetases, multiple proteins involved in the assembly of electron transport chain complexes, and several mitochondrial carrier proteins. While this article focuses on the ctenophore *M. leidyi*, MitoPredictor can be useful for predicting and analyzing mitochondrial proteomes of other animal species.

## References

1. Hatefi Y. The mitochondrial electron transport and oxidative phosphorylation system. Annu Rev Biochem. 1985;54:1015–1069. doi:10.1146/annurev.bi.54.070185.005055

2. Stehling O, Wilbrecht C, Lill R. Mitochondrial iron–sulfur protein biogenesis and human disease. Biochimie. 2014;100:61–77. doi:10.1016/j.biochi.2014.01.010

3. Guda P, Guda C, Subramaniam S. Reconstruction of Pathways Associated with Amino Acid Metabolism in Human Mitochondria. Genomics, Proteomics & Bioinformatics. 2007;5(3):166–176. doi:10.1016/S1672-0229(08)60004-2

4. Mayr JA. Lipid metabolism in mitochondrial membranes. Journal of Inherited Metabolic Disease. 2015;38(1):137–144. doi:10.1007/s10545-014-9748-x

5. Oberst A, Bender C, Green DR. Living with death: the evolution of the mitochondrial pathway of apoptosis in animals. Cell Death & Differentiation. 2008;15(7):1139–1146. doi:10.1038/cdd.2008.65

6. Wang C, Youle RJ. The Role of Mitochondria in Apoptosis. Annu Rev Genet. 2009;43(1):95–118. doi:10.1146/annurev-genet-102108-134850

7. Chandel NS. Mitochondria as signaling organelles. BMC Biology. 2014;12(1):34. doi:10.1186/1741-7007-12-34

8. Meisinger C, Sickmann A, Pfanner N. The mitochondrial proteome: from inventory to function. Cell. 2008;134(1):22–24.

9. Wiedemann N, Pfanner N. Mitochondrial Machineries for Protein Import and Assembly. Annu Rev Biochem. 2017;86(1):685–714. doi:10.1146/annurev-biochem-060815-014352

10. Calvo SE, Clauser KR, Mootha VK. MitoCarta2.0: an updated inventory of mammalian mitochondrial proteins. Nucleic Acids Research. 2016;44(D1):D1251–D1257. doi:10.1093/nar/gkv1003

11. Smith AC, Robinson AJ. MitoMiner v3.1, an update on the mitochondrial proteomics database. Nucleic Acids Research. 2016;44(D1):D1258–D1261. doi:10.1093/nar/gkv1001

12. Li J, Cai T, Wu P, et al. Proteomic analysis of mitochondria from Caenorhabditis elegans. Proteomics. 2009;9(19):4539–4553.

13. Hu Y, Comjean A, Perkins LA, Perrimon N, Mohr SE. GLAD: an Online Database of Gene List Annotation for Drosophila. J Genomics. 2015;3:75–81. doi:10.7150/jgen.12863

14. Heazlewood JL, Howell KA, Whelan J, Millar AH. Towards an Analysis of the Rice Mitochondrial Proteome. Plant Physiology. 2003;132(1):230–242. doi:10.1104/pp.102.018986

15. Salvato F, Havelund JF, Chen M, et al. The Potato Tuber Mitochondrial Proteome. Plant Physiology. 2014;164(2):637–653. doi:10.1104/pp.113.229054

16. Millar AH, Sweetlove LJ, Giegé P, Leaver CJ. Analysis of the Arabidopsis Mitochondrial Proteome. Plant Physiology. 2001;127(4):1711–1727. doi:10.1104/pp.010387

17. Rao RSP, Salvato F, Thal B, Eubel H, Thelen JJ, Møller IM. The proteome of higher plant mitochondria. Mitochondrion. 2017;33:22–37. doi:10.1016/j.mito.2016.07.002

18. Cherry JM, Hong EL, Amundsen C, et al. Saccharomyces Genome Database: the genomics resource of budding yeast. Nucleic Acids Research. 2012;40(D1):D700–D705. doi:10.1093/nar/gkr1029

19. Gawryluk RMR, Chisholm KA, Pinto DM, Gray MW. Compositional complexity of the mitochondrial proteome of a unicellular eukaryote (Acanthamoeba castellanii, supergroup Amoebozoa) rivals that of animals, fungi, and plants. Journal of Proteomics. 2014;109:400–416. doi:10.1016/j.jprot.2014.07.005

20. Lavrov DV, Pett W. Animal Mitochondrial DNA as We Do Not Know It: mt-Genome Organization and Evolution in Nonbilaterian Lineages. Genome Biology and Evolution. 2016;8(9):2896–2913. doi:10.1093/gbe/evw195

21. Emms DM, Kelly S. OrthoFinder: phylogenetic orthology inference for comparative genomics. Genome Biology. 2019;20(1):238. doi:10.1186/s13059-019-1832-y

22. Persson E, Sonnhammer ELL. InParanoid-DIAMOND: faster orthology analysis with the InParanoid algorithm. Bioinformatics. 2022;38(10):2918–2919. doi:10.1093/bioinformatics/btac194

23. Lechner M, Findeiß S, Steiner L, Marz M, Stadler PF, Prohaska SJ. Proteinortho: Detection of (Co-)orthologs in large-scale analysis. BMC Bioinformatics. 2011;12(1):124. doi:10.1186/1471-2105-12-124

24. Emanuelsson O, Nielsen H, Brunak S, von Heijne G. Predicting Subcellular Localization of Proteins Based on their N-terminal Amino Acid Sequence. Journal of Molecular Biology. 2000;300(4):1005–1016. doi:10.1006/jmbi.2000.3903

25. Fukasawa Y, Tsuji J, Fu SC, Tomii K, Horton P, Imai K. MitoFates: Improved Prediction of Mitochondrial Targeting Sequences and Their Cleavage Sites *[S]. Molecular & Cellular Proteomics. 2015;14(4):1113–1126. doi:10.1074/mcp.M114.043083

26. Yu CS, Chen YC, Lu CH, Hwang JK. Prediction of protein subcellular localization. Proteins: Structure, Function, and Bioinformatics. 2006;64(3):643–651. doi:10.1002/prot.21018

27. King BR, Guda C. ngLOC: an n-gram-based Bayesian method for estimating the subcellular proteomes of eukaryotes. Genome Biology. 2007;8(5):R68. doi:10.1186/gb-2007-8-5-r68

28. Kumar R, Kumari B, Kumar M. Proteome-wide prediction and annotation of mitochondrial and sub-mitochondrial proteins by incorporating domain information. Mitochondrion. 2018;42:11–22. doi:10.1016/j.mito.2017.10.004

29. Muthye V, Kandoi G, Lavrov DV. MMPdb and MitoPredictor: Tools for facilitating comparative analysis of animal mitochondrial proteomes. Mitochondrion. 2020;51:118–125. doi:10.1016/j.mito.2020.01.001

30. Salvatore M, Warholm P, Shu N, Basile W, Elofsson A. SubCons: a new ensemble method for improved human subcellular localization predictions. Bioinformatics. 2017;33(16):2464–2470. doi:10.1093/bioinformatics/btx219

31. Briesemeister S, Blum T, Brady S, Lam Y, Kohlbacher O, Shatkay H. SherLoc2: A High-Accuracy Hybrid Method for Predicting Subcellular Localization of Proteins. J Proteome Res. 2009;8(11):5363–5366. doi:10.1021/pr900665y

32. Goldberg T, Hamp T, Rost B. LocTree2 predicts localization for all domains of life. Bioinformatics. 2012;28(18):i458–i465. doi:10.1093/bioinformatics/bts390

33. Blum T, Briesemeister S, Kohlbacher O. MultiLoc2: integrating phylogeny and Gene Ontology terms improves subcellular protein localization prediction. BMC Bioinformatics. 2009;10(1):274. doi:10.1186/1471-2105-10-274

34. Schultz DT, Haddock SHD, Bredeson JV, Green RE, Simakov O, Rokhsar DS. Ancient gene linkages support ctenophores as sister to other animals. Nature. 2023;618(7963):110–117. doi:10.1038/s41586-023-05936-6

35. Pett W, Ryan JF, Pang K, et al. Extreme mitochondrial evolution in the ctenophore Mnemiopsis leidyi: Insight from mtDNA and the nuclear genome. Mitochondrial DNA. 2011;22(4):130–142. doi:10.3109/19401736.2011.624611

36. Ryan JF, Pang K, Schnitzler CE, et al. The Genome of the Ctenophore Mnemiopsis leidyi and Its Implications for Cell Type Evolution. Science. 2013;342(6164):1242592. doi:10.1126/science.1242592

37. Moreland RT, Nguyen AD, Ryan JF, et al. A customized Web portal for the genome of the ctenophore Mnemiopsis leidyi. BMC Genomics. 2014;15(1):316. doi:10.1186/1471-2164-15-316

38. Moroz LL, Kocot KM, Citarella MR, et al. The ctenophore genome and the evolutionary origins of neural systems. Nature. 2014;510(7503):109–114. doi:10.1038/nature13400

39. Johnson SB, Winnikoff JR, Schultz DT, et al. Speciation of pelagic zooplankton: Invisible boundaries can drive isolation of oceanic ctenophores. Front Genet. 2022;13:970314. doi:10.3389/fgene.2022.970314

40. Schultz DT, Francis WR, McBroome JD, Christianson LM, Haddock SHD, Green RE. A chromosome-scale genome assembly and karyotype of the ctenophore Hormiphora californensis. G3 (Bethesda). 2021;11(11). doi:10.1093/g3journal/jkab302

41. Schultz DT, Eizenga JM, Corbett-Detig RB, Francis WR, Christianson LM, Haddock SHD. Conserved novel ORFs in the mitochondrial genome of the ctenophore Beroe forskalii. PeerJ. 2020;8:e8356. doi:10.7717/peerj.8356

42. Kohn AB, Citarella MR, Kocot KM, Bobkova YV, Halanych KM, Moroz LL. Rapid evolution of the compact and unusual mitochondrial genome in the ctenophore, Pleurobrachia bachei. Mol Phylogenet Evol. 2012;63(1):203–207. doi:10.1016/j.ympev.2011.12.009

43. Wang M, Cheng F. The complete mitochondrial genome of the Ctenophore Beroe cucumis, a mitochondrial genome showing rapid evolutionary rates. Mitochondrial DNA B Resour. 2019;4(2):3774–3775. doi:10.1080/23802359.2019.1580165

44. Fu L, Niu B, Zhu Z, Wu S, Li W. CD-HIT: accelerated for clustering the next-generation sequencing data. Bioinformatics. 2012;28(23):3150–3152. doi:10.1093/bioinformatics/bts565

45. Koskinen P, Törönen P, Nokso-Koivisto J, Holm L. PANNZER: high-throughput functional annotation of uncharacterized proteins in an error-prone environment. Bioinformatics. 2015;31(10):1544–1552. doi:10.1093/bioinformatics/btu851

46. Fang H, Gough J. dcGO: database of domain-centric ontologies on functions, phenotypes, diseases and more. Nucleic Acids Research. 2013;41(D1):D536–D544. doi:10.1093/nar/gks1080

47. Herwig R, Hardt C, Lienhard M, Kamburov A. Analyzing and interpreting genome data at the network level with ConsensusPathDB. Nature Protocols. 2016;11(10):1889–1907. doi:10.1038/nprot.2016.117

48. Mi H, Muruganujan A, Ebert D, Huang X, Thomas PD. PANTHER version 14: more genomes, a new PANTHER GO-slim and improvements in enrichment analysis tools. Nucleic Acids Research. 2019;47(D1):D419–D426. doi:10.1093/nar/gky1038

49. Mi H, Muruganujan A, Casagrande JT, Thomas PD. Large-scale gene function analysis with the PANTHER classification system. Nature Protocols. 2013;8(8):1551–1566. doi:10.1038/nprot.2013.092

50. Szklarczyk D, Gable AL, Lyon D, et al. STRING v11: protein–protein association networks with increased coverage, supporting functional discovery in genome-wide experimental datasets. Nucleic Acids Research. 2019;47(D1):D607–D613. doi:10.1093/nar/gky1131

51. Harris TW, Arnaboldi V, Cain S, et al. WormBase: a modern Model Organism Information Resource. Nucleic Acids Research. 2020;48(D1):D762–D767. doi:10.1093/nar/gkz920

52. Lyne R, Smith R, Rutherford K, et al. FlyMine: an integrated database for Drosophila and Anopheles genomics. Genome Biology. 2007;8(7):R129. doi:10.1186/gb-2007-8-7-r129

53. Pett W, Lavrov DV. Cytonuclear Interactions in the Evolution of Animal Mitochondrial tRNA Metabolism. Genome Biology and Evolution. 2015;7(8):2089–2101. doi:10.1093/gbe/evv124

54. Palmieri F. Mitochondrial carrier proteins. FEBS Letters. 1994;346(1):48–54. doi:10.1016/0014-5793(94)00329-7

55. Palmieri F, Pierri CL, De Grassi A, Nunes-Nesi A, Fernie AR. Evolution, structure and function of mitochondrial carriers: a review with new insights. The Plant Journal. 2011;66(1):161–181. doi:10.1111/j.1365-313X.2011.04516.x

56. Pao Stephanie S., Paulsen Ian T., Saier Milton H. Major Facilitator Superfamily. Microbiology and Molecular Biology Reviews. 1998;62(1):1–34. doi:10.1128/mmbr.62.1.1-34.1998

57. Yan N. Structural Biology of the Major Facilitator Superfamily Transporters. Annu Rev Biophys. 2015;44(1):257–283. doi:10.1146/annurev-biophys-060414-033901

